# Magnetic Levitation of Intact Tumor Biopsies Reveals Multivariate Biophysical Signatures of Breast Cancer Aggressiveness

**DOI:** 10.64898/2026.06.26.734698

**Authors:** Meltem Guzelgulgen, Zehra Elif Gunyuz, Muge Anil-Inevi, Devrim Pesen-Okvur, Betul Bolat-Kucukzeybek, Merve Gursoy, Ozden Yalcin-Ozuysal, Gulistan Mese, Engin Ozcivici

**Affiliations:** Department of Bioengineering, Izmir Institute of Technology, Izmir, Turkey; Department of Molecular Biology and Genetics, Izmir Institute of Technology, Izmir, Turkey; Department of Pathology, Izmir Katip Celebi University Ataturk Training and Research Hospital, Izmir, Turkey; Department of Radiology, Acibadem Izmir Kent Hospital, Breast Clinic, Izmir, Turkey

**Keywords:** Breast cancer, magnetic levitation, tissue biopsy, biophysical profiling, tumor heterogenity, latent variable analysis

## Abstract

Diagnostic assessment of breast cancer biopsies remains reliant on resource-intensive histopathology and molecular profiling, which often lack real-time physiological readouts. Magnetic levitation (MagLev) enables label-free density profiling of single cells, yet its application to intact tissue biopsies has been precluded by size-dependent geometric artifacts and the absence of analytical frameworks for biopsy-scale samples. Here, we report the first application of MagLev to intact invasive breast carcinoma biopsies (200-600 μm) for biophysical profiling, generating multivariate biophysical signatures from 203 samples across 17 patients. We developed a physics-based size-correction algorithm (xmc) that isolates biological density from geometric artifact, and demonstrate that tissue viability is predicted not by average levitation height, but by spatial heterogeneity across replicate samples, reflecting the microenvironmental complexity of metabolically active tumors. Multivariate integration using Partial Least Squares (PLS) regression and Factor Analysis of Mixed Data (FAMD) identified nodal status (N) as the strongest biophysical predictor, suggesting that lymphatic dissemination capacity leaves a measurable signature in the primary tumor density profile. Unsupervised patient clustering in PLS-derived latent space recovered three clinically coherent subgroups aligned with molecular subtypes. This 30-minute, low-cost assay provides exploratory biophysical stratification complementary to existing diagnostics, particularly in resource-limited settings.

## INTRODUCTION

Breast cancer is the most commonly diagnosed malignancy in women globally, accounting for 11.6% of new cancer cases and 6.9% of cancer-related deaths, with 2.3 million diagnoses and 670,000 deaths reported in 2022 (1,2). Current clinical management relies on molecular markers—estrogen receptor (ER), progesterone receptor (PR), and HER2—to guide therapeutic decisions. However, intratumoral heterogeneity fundamentally limits the predictive power of single-biopsy molecular profiling (3,4). Regional HER2 expression variability occurs in up to 40% of tumors and correlates with anti-HER2 resistance and reduced survival (4), while architectural and metabolic heterogeneity—reflecting hypoxic gradients, necrotic foci, and variable stromal infiltration—contributes to relapse in 20% of early-stage patients despite curative-intent treatment (3). Existing strategies to address this challenge, including PET/CT metabolic profiling (5), patient-derived organoids, liquid biopsies, and quantitative MRI heterogeneity indices (6), remain resource-intensive and infrastructure-dependent, limiting accessibility in the low- and middle-income settings where the majority of the global breast cancer burden occurs.

Physical phenotyping provides a complementary approach to molecular diagnostics, exploiting measurable biophysical properties—density, stiffness, electrical impedance, and morphology—that reflect tumor cell state and microenvironmental architecture (7–9). Unlike genomic or proteomic profiling, biophysical measurements provide rapid, label-free characterization that integrates metabolic activity (affecting density through osmotic regulation and organelle distribution), structural organization (cellularity, extracellular matrix composition, stromal content), and functional state (proliferation, differentiation) (10–14). Atomic force microscopy has quantified stiffness differences between benign and malignant breast tissue (15,16), deformability cytometry discriminates metastatic from non-metastatic circulating tumor cells (17), and radiomics extracts quantitative texture features from CT and MRI that predict treatment response and survival independently of molecular subtype (18). These approaches share key translational advantages involving minimal sample preparation, rapid turnaround, and the ability to detect phenotypic variation invisible to genomic classifiers—cells with identical genotypes can exhibit distinct density or mechanical phenotypes depending on microenvironmental context or metabolic programming (19,20).

Magnetic levitation (MagLev) exploits density-dependent equilibrium positioning in paramagnetic media within magnetic field gradients to achieve label-free separation of biological samples (21,22). In a two-magnet Anti-Helmholtz configuration, diamagnetic objects suspended in paramagnetic medium levitate at heights inversely related to their density, with sensitivity approaching 0.001 g/cm³ (23). Single-cell MagLev studies have established that viable, apoptotic, and necrotic cells occupy distinct levitation zones (24), enabling applications in drug response profiling (25) and circulating tumor cell isolation (26). The technology has also been extended to three-dimensional tissue engineering, where density-driven self-assembly generates scaffold-free constructs (27,28). Despite this progress, no previous study has applied MagLev to intact clinical tissue biopsies. Tissue samples are 20–30-fold larger than single cells (200–600 μm vs. 10–20 μm), incorporate extracellular matrix, stromal components, and spatial architecture absent from dissociated suspensions, and span 10–40% of the inter-magnet gap—causing size-dependent geometric artifacts that confound density measurements derived from single-cell analytical frameworks. Overcoming these challenges is essential to translate MagLev’s speed and accessibility to clinical specimens.

Here, we present the first application of magnetic levitation to intact tissue biopsies, developing a multivariate biophysical profiling platform for invasive breast carcinoma. Analyzing 203 samples from 17 patients, we also introduced physics-based size correction and statistical residualization on levitation metrics to isolate biophysical signals from geometric artifacts related to punch biopsies. We demonstrate that sample based spatial heterogeneity in levitation profiles, but not average density, predicts tissue viability, inverting the single-cell paradigm and revealing that architectural complexity encodes biological information. Multivariate integration identifies two orthogonal biological axes: a viability-burden dimension dominated by metabolic state and nodal involvement, and an architectural dimension reflecting histological subtype and tumor size. Nodal status emerges as the strongest biophysical predictor, and unsupervised patient projection into this latent space recovers three naturally segregating clusters aligned with molecular subtypes. This work establishes tissue-scale MagLev as a rapid, label-free platform for clinically relevant patient stratification, with particular value in resource-limited settings.

## MATERIALS AND METHODS

### Patient Sample Collection and Ethical Approval

This study was conducted under the approval of the İzmir Katip Çelebi University Faculty of Medicine Clinical Research Ethics Committee (Approval Number: E-32623740-604.01.01-2300046144, Decision Number: 2023/12). Tumor tissue samples were obtained from 17 patients diagnosed with invasive breast carcinoma with informed consent between 03/2024 – 07/2025 at the İzmir Katip Çelebi University Atatürk Training and Research Hospital Radiology Clinic. Core needle biopsy procedure was performed on patients with malignant breast mass under local anesthesia using a 14G needle. Clinical and pathological data were extracted from medical records, including patient age, primary tumor size (T stage), regional lymph node involvement (N stage), distant metastasis (M stage), histological subtype (invasive ductal carcinoma [IDC], invasive lobular carcinoma [ILC], invasive tubular carcinoma [ITC], neuroendocrine [NE]), histological and nuclear grade (1–3), receptor status (ER, PR, HER2), Ki67 proliferation index (%), tumor diameter (mm), and lymph node count. Molecular subtypes were classified as Luminal A (LumA), Luminal B (LumB), HER2-enriched, and triple-negative breast cancer (TNBC) based on the IHC surrogate definitions established by the St. Gallen International Breast Cancer Consensus. A composite Aggressiveness Score (0–10 scale) to function as a weighted cumulative prognostic index was calculated as: (Ki67≥14%) + (Nuclear Grade=3) + (Histological Grade=3) + (T≥3) + (N≥1) + (M=1) + (Lymph Nodes≥4) + (Molecular=TNBC) + (PET=1) + 0.5×(HER2=1), with scores dichotomized as non-aggressive (≤4) versus aggressive (>4) for categorical analyses (29).

### Tissue Biopsy Preparation and Processing

Approximately 1 cm long biopsy specimens obtained from patients were brought to the laboratory in a serum-free DMEM/F12 nutrient medium with 1% Pen/Strep, maintaining a cold chain. Using sequential razor blade sectioning perpendicular to the tissue’s long axis, specimens were divided into 400 μm-thick cylindrical discs and maintained in MCF10A complete medium (5% horse serum, 20 ng/mL EGF, 0.5 mg/mL hydrocortisone, 100 ng/mL cholera toxin, 10 μg/mL insulin, 1% Pen/Strep in DMEM/F12) at 37°C with 5% CO₂ throughout the experiment.

For viability assessment, patient-matched tissue slices cultured in MCF10A complete medium, 37°C and 5% CO_2_ for 1-2 hours prior to viability assay to stabilize metabolic activity post-resection. For magnetic levitation experiments, samples were extracted from tissue slices using blunt-tipped 22-gauge needles. Samples were taken from both peripheral and core regions to capture intra-tumoral heterogeneity. Sample cross-sectional dimensions were confirmed to fall within 200–600 μm by brightfield microscopy prior to inclusion. Samples were maintained in MCF10A complete medium, 37°C and 5% CO_2_ for 2-6 hours prior to magnetic levitation.

### Tissue Viability Measurements

Tissue viability was quantified using PrestoBlue™ Cell Viability Reagent (Invitrogen, A13261), a resazurin-based metabolic assay reporting mitochondrial reducing capacity. Reagent was diluted 1:10 (v/v) in culture medium and applied for 90 min at 37°C in a humidified 5% CO₂ incubator. Absorbance was measured at 570 nm (experimental) and 600 nm (reference) using a microplate spectrophotometer (Thermo Scientific Multiskan SkyHigh). Percent reagent reduction was calculated using molar extinction coefficients of the oxidized and reduced forms, correcting for media background and spectral overlap. Two patient-level viability metrics were derived: mean metabolic activity across replicate slices (Viability_avg) and its standard deviation (Viability_stdev), reflecting intra-tumoral metabolic heterogeneity. The cytotoxicity of gadolinium (Gadobutrol; Gadavist, Bayer, Germany) at concentrations up to 200 mM was assessed in parallel to confirm biocompatibility of the levitation medium. Only patients passing the viability acceptance criterion were included in the study.

### Magnetic Levitation Setup, Image Acquisition

Magnetic levitation was performed with a custom system comprising two neodymium permanent magnets (NdFeB, N52 grade; 50×2×5 mm; Supermagnete) in a repulsive Anti- Helmholtz configuration with ∼1.5 mm vertical separation, mounted in a photoreactive resin holder fabricated by 3D printing (Formlabs Form 2). A glass capillary (1×1 mm cross-section, 50 mm length; Vitrocom) was positioned vertically between the magnets along the central axis. Tilted side mirrors (45°) enabled visualization of levitated samples using an inverted microscope (Olympus IX-83). The magnetic midpoint (x=0 μm) was defined as the equidistant plane between the two magnet faces. Paramagnetic levitation medium was prepared by dissolving gadobutrol (Gadavist, 150 mM) in MCF10A complete medium. Individual biopsy puncture samples were gently loaded into the capillary (50 μL levitation medium) by micropipette, avoiding air bubbles and tissue damage. Samples equilibrated for 2–3 minutes before imaging at 4× magnification. A calibration scale (116 pixels = 200 μm) was imaged under identical conditions for pixel-to-micrometer conversion.

### Image Processing and Levitation Metrics

Levitation images were processed in ImageJ Fiji using a pipeline of thresholding, edge detection, and contour segmentation to define tissue boundaries. For each segmented sample the following morphological parameters were extracted: Area (μm²), Perimeter (μm), Height (maximum vertical span, μm), Width (maximum horizontal span, μm), and Circularity (4π×Area/Perimeter²; 1.0 = perfect circle). Levitation heights were measured relative to the magnetic midpoint (Supplementary Figure S1). Three raw metrics were calculated: (1) xm, the sample center-of-gravity height; (2) xmin, the minimum levitation height (bottom tissue surface); and (3) xmax, the maximum levitation height (top tissue surface). Negative values denote positions below the magnetic midpoint; positive values denote positions above. Furthermore, to isolate density-related signals from geometric artifacts, three corrected metrics were derived. The physics-based correction (xmc) recalculates the center-of-gravity after computationally excluding all tissue positioned above the magnetic midpoint (x≥0 μm), under the assumption that above-midpoint tissue represents geometric extension rather than density-driven positioning (28) with the formulation: if xmax<midpoint, xmc = xm; else xmc= (midpoint-(xmax-midpoint)+xmin)/2 (Supplementary Figure S1). Statistical residualization (xm_res, xmin_res) regressed xm and xmin against Area across all samples (n=203) using ordinary least squares, retaining residuals as size-independent metrics that by construction show zero correlation with Area. Division-based normalization (xm/Area, etc.) was tested but paradoxically increased size correlations due to spurious correlation effects (30) and was excluded from further analysis (Supplementary Figure S2).

Patient-level summary features were derived by aggregating per-sample metrics: six averages (central tendency of density properties across replicate samples) and six heterogeneity measures (standard deviation across replicate samples), the latter quantifying spatial and sampling variability within each tumor.

### Statistical Analysis

All analyses were performed in Python 3 using NumPy, SciPy, pandas, scikit-learn, statsmodels, Matplotlib, and Seaborn libraries. Statistical significance was defined as p<0.05; statistical trend as p<0.20. Pearson correlation coefficients (r) assessed linear associations between continuous variables; Spearman rank correlations (ρ) were used for ordinal variables. Central tendency of indices was measured with average values while standard deviations were used as heterogeneity measures. Associations between levitation metrics and categorical clinical variables (molecular subtype, histological subtype, receptor status, TNM stage, grade) were assessed by two-sample t-tests (binary variables; effect size: Cohen’s d), one-way ANOVA (multi-level categorical variables; effect size: η²) with Tukey HSD post-hoc testing, and Welch’s ANOVA where Levene’s test indicated heterogeneity of variance. Multiple linear regression with stepwise backward elimination (AIC-optimized) was performed with all 12 levitation features as predictors and Viability_avg as the outcome (n=17). Full details of FAMD, PLS-R, factor analysis, and latent space analysis methods are provided in Supplementary Methods.

## RESULTS

### Tissue Levitation Profiles and Size-Dependent Artifacts

We measured magnetic levitation profiles for 203 invasive breast carcinoma biopsies from 17 patients (mean 12 samples per patient, range 4–21), quantifying center-of-gravity (xm), minimum surface (xmin), and maximum surface (xmax) levitation heights alongside morphological parameters (Figure 1A; Supplementary Figure S1; Supplementary Tables S1–S2). Samples showed substantial dimensional variability (Area: 208000±59000 μm²; Height: 499±77 μm; Width: 622±132 μm), reflecting size heterogeneity in punch biopsies. Raw levitation metrics varied correspondingly: xm averaged −150±77 μm (range: −327 to +10 μm), xmin averaged −399±81 μm (range: −574 to −227 μm), while xmax averaged +100±91 μm (range: −164 to +260 μm). Notably, xmax was predominantly positive, indicating most samples extended above the magnetic midpoint—an unexpected finding given that breast carcinoma tissue density (>1.04 g/cm³ (31)) should position samples entirely below the midpoint in our paramagnetic medium.

**Figure 1.**
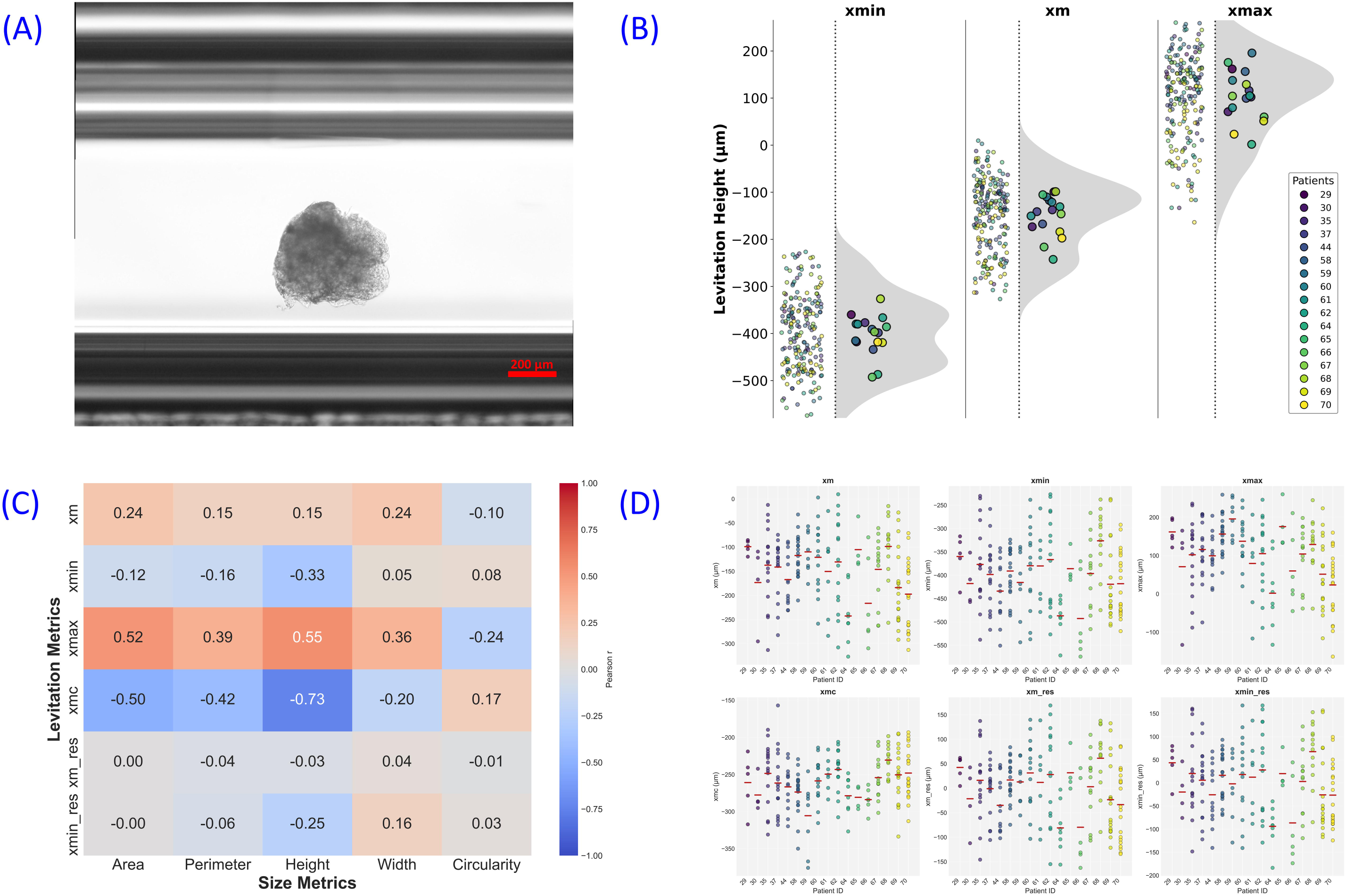
Tissue levitation platform and size-correction framework. (A) Schematic of the magnetic levitation system showing the anti-Helmholtz magnet configuration, glass capillary, and tilted side mirrors enabling brightfield visualization of levitated tissue biopsies. Scale bar: 200 μm. (B) Raw levitation metrics xmin, xm, and xmax distributions on biopsy sample (n=203) and patient level (n=17). Color distribution represents patients. (C) Correlation matrix (Spearman ρ) between the six levitation metrics, demonstrating that physics-based correction (xmc) and statistical residualization (xm_res, xmin_res) capture largely orthogonal information from each other and from raw metrics. (D) Schematic summary of the six levitation metrics retained for downstream analysis: three raw (xm, xmin, xmax) and three corrected (xmc, xm_res, xmin_res), each representing complementary aspects of tissue biophysics.

We attributed this observation to a size-dependent geometric artifact unique to biopsy-scale levitation: tissue dimensions (200–600 μm) span 10–40% of the inter-magnet gap (∼1.5 mm), causing different tissue regions to experience substantially different field strengths rather than behaving as point masses. Correlation analysis confirmed strong size-dependency for xmax (Area: r=0.52, p<0.001; Height: r=0.55, p<0.001; Width: r=0.36, p<0.001), with weaker but significant associations for xm and xmin (Supplementary Table S3). Physics-based correction (xmc) eliminated positive values entirely (xmc mean: −260.48 μm), while statistical residualization yielded xm_res and xmin_res with zero correlation to Area by construction (Supplementary Figure S2). The two correction approaches captured orthogonal information (xmc vs. xm_res: ρ=0.40; xmc vs. xmin_res: ρ=0.52; Figure 1C), motivating retention of all 12 summary features: six averages (central tendency) and six heterogeneity measures (standard deviation across replicate samples), for downstream analysis (Figure 1D).

### Spatial Heterogeneity Predicts Tissue Viability

Tissue viability measured by PrestoBlue metabolic assay (n=17 patients) showed no significant correlations between patient-averaged levitation metrics and mean metabolic activity (Viability_avg; strongest association: xmc_avg vs. Viability_avg: r=0.30, p=0.24; Supplementary Table S4, Supplementary Figure S3). This contrasts with single-cell levitation studies, where viable cells consistently levitate higher than apoptotic or necrotic cells due to membrane permeabilization and cytoplasmic condensation (24,25). Size-corrected residual metrics similarly yielded null results, indicating that size-normalization alone does not reveal viability–density relationships at the tissue scale.

In contrast, levitation heterogeneity (the sample-to-sample standard deviation within each patient) revealed the only significant univariate viability association. Heterogeneity_xmc (SD of xmc across ∼12 replicate samples per patient) correlated strongly with mean viability (r=0.57, p=0.017; Figure 2; Supplementary Table S4). This relationship was specific to the physics-corrected metric; raw heterogeneity measures showed weaker or null associations (Heterogeneity_xm: r=−0.05, p=0.86; Heterogeneity_xmax: r=−0.26, p=0.31). Statistical trends were also observed for heterogeneity of residual metrics versus viability dispersion (Heterogeneity_xm_res vs. Viability_stdev: r=0.47, p=0.059; Supplementary Table S5), suggesting that spatial heterogeneity in both levitation and metabolism co-vary at the tissue level. Multivariate linear regression incorporating all levitation features jointly explained 76% of viability variance (adjusted R²=0.76; F-test p=0.019; Supplementary Table S6), with Heterogeneity_xmc remaining a significant independent contributor (β=0.59, p=0.009).

**Figure 2.**
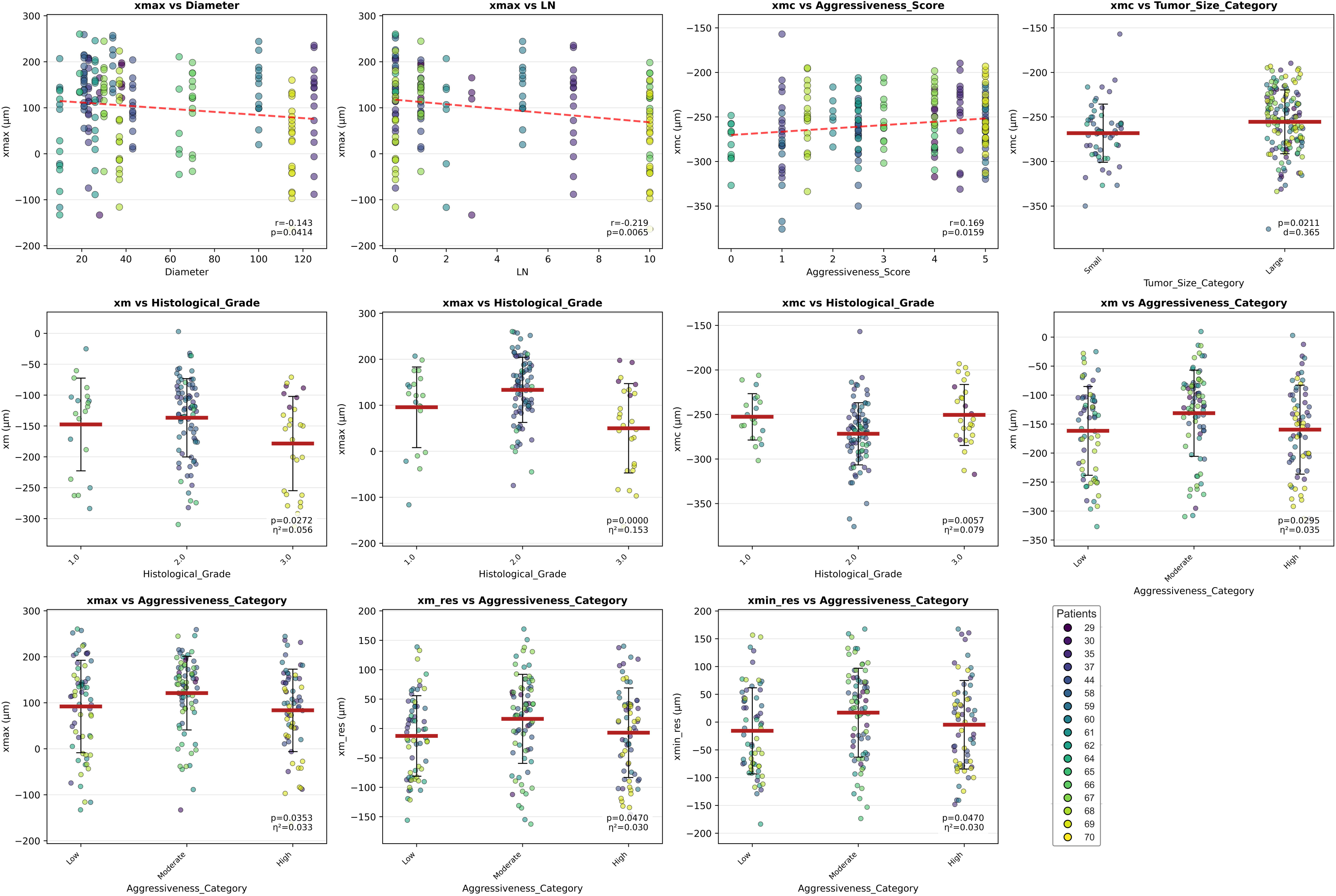
Clinical associations of levitation metrics at the sample level (n = 203 biopsies). Significant associations (p < 0.05) between levitation metrics and clinical variables identified from 72 pairwise comparisons (6 metrics × 12 variables). The first three panels show scatter plots for significant continuous associations; Pearson r and p-values are annotated. The remaining eight panels depict categorical associations, each showing individual sample distributions with group means (red horizontal lines) and SD whiskers; significant group differences (p < 0.05) are annotated with effect size (Cohen’s d for binary comparisons, η² for ANOVA). Color distribution represents patients.

### Clinical Associations of Levitation Metrics

Given substantial intra-tumoral heterogeneity in levitation profiles (mean intraclass correlation coefficient ICC=0.17, indicating only 17% of variance resides between patients; Supplementary Table S7), we treated individual sample measurements (n=203) as the primary analytical unit, capturing spatial variability as biological signal rather than technical noise. From 72 comparisons (6 metrics × 12 clinical variables), 11 associations reached significance (Figure 2).

The raw metric xmax showed the strongest and most numerous clinical correlations: negative associations with tumor diameter (r=−0.14, p=0.041), lymph node involvement (r=−0.22, p=0.007), and histological grade (ANOVA: F=11.3, p<0.001, η²=0.15; grade 3 tumors showed xmax 40 μm lower than grade 1), as well as discrimination of molecular subtypes (F=3.2, p=0.025) and T-stage (Cohen’s d=0.31, p=0.047). The physics-corrected metric xmc revealed complementary and directionally opposite patterns: positive correlation with Aggressiveness Score (r=0.17, p=0.016), discrimination of tumor size category (Cohen’s d=0.37, p=0.021), and association with histological grade (F=5.4, p=0.006, η²=0.08). This directional divergence—where aggressive tumors levitate lower by xmax but higher by xmc—suggests that vertical density stratification within biopsies encodes biological information: aggressive tumors may exhibit distinct density gradients compared to less aggressive tumors with more uniform internal density distributions. Statistical residualization metrics (xm_res, xmin_res) showed weaker but consistent associations with Aggressiveness Category (both p=0.047, η²=0.03), confirming size-independent density differences at smaller effect sizes.

Complementary patient-level analysis (n=17) using a relaxed significance threshold (p<0.20 to detect trends given limited sample size) identified additional associations (Supplementary Figure S4). Mean viability showed the strongest patient-level associations: Aggressiveness Category (ANOVA: F=14.4, p=0.0004, η²=0.67; aggressive tumors had 85% higher viability) and Nuclear Grade (Cohen’s d=−2.01, p=0.004; grade 3 showed 73% higher viability than grade 1-2). Patient-level heterogeneity metrics revealed trends with molecular subtype (Heterogeneity_xmin: F=3.5, p=0.034) and tumor size (Heterogeneity_xmin vs. Diameter: r=0.48, p=0.050), reinforcing that levitation heterogeneity captures clinically relevant biological complexity. The convergence of sample-level and patient-level findings—despite analyzing different units of observation (individual biopsies vs. patient means)—provides orthogonal validation that levitation metrics encode genuine biological signals robust to analytical approach.

### Multivariate Structure of Levitation Profiles

Factor Analysis of Mixed Data (FAMD) integrating 8 levitation metrics, 2 viability measures, and 8 clinical variables (n=17 patients) reduced the 18-dimensional variable space into 10 components explaining 94.9% of total variance (Figure 3A; Supplementary Figures S5–S6). PC1 (21.2% variance) loaded positively on raw levitation averages (xm, xmin, xmax: loadings 0.29–0.31) and inversely on heterogeneity metrics, reflecting a fundamental trade-off between mean tissue density and spatial variability. PC2 (20.0% variance) captured a tissue complexity dimension loaded by viability (0.32), tumor diameter (0.31), Aggressiveness Score (0.31), and xmc_avg (0.25), contrasted by patient age (−0.18)—separating metabolically active, aggressive tumors from smaller, lower-viability cases.

**Figure 3.**
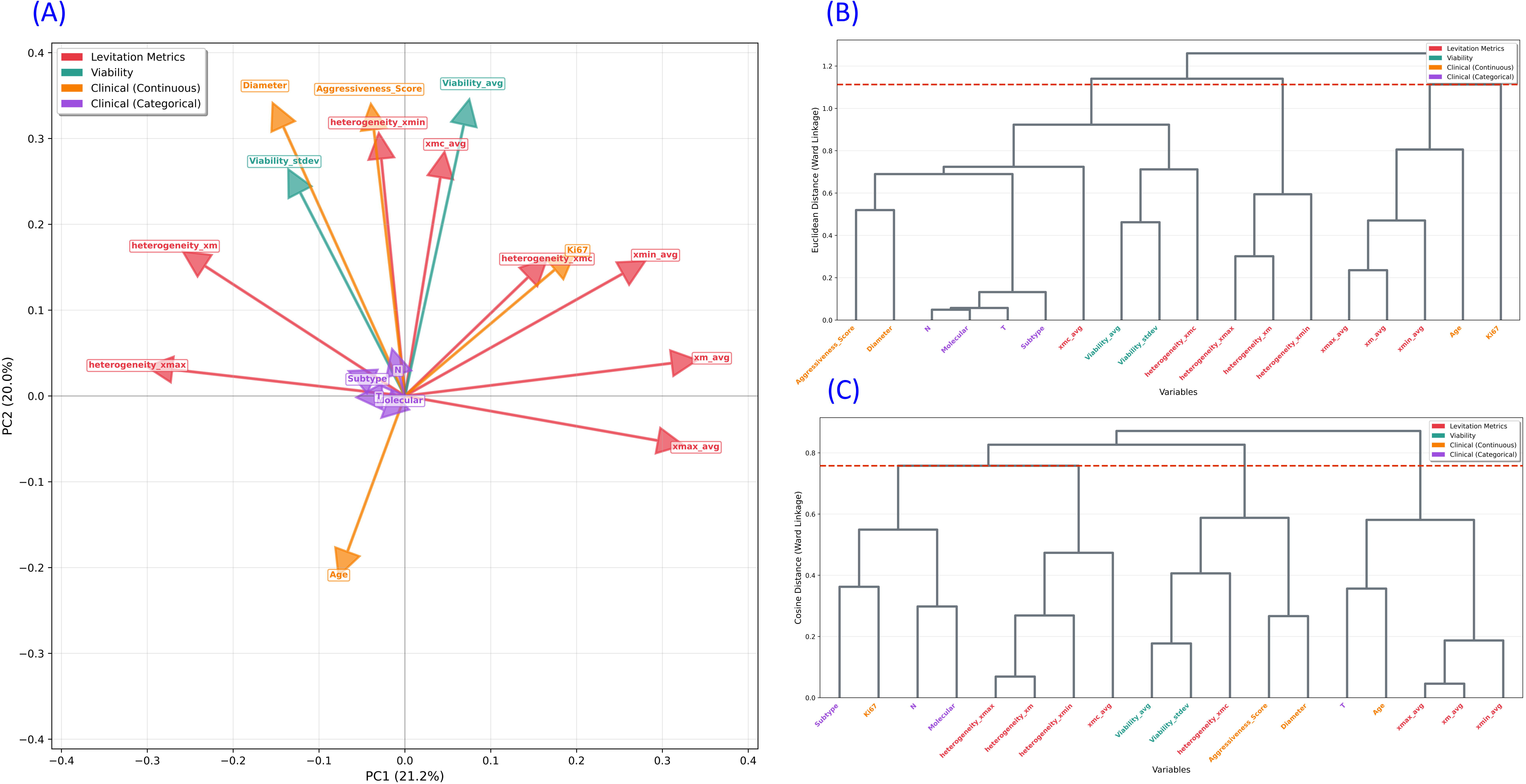
Multivariate structure revealed by Factor Analysis of Mixed Data. (A) FAMD biplot showing variable projections onto PC1 and PC2. Levitation metrics are shown in blue, viability measures in green, and clinical variables in orange and purple. PC1 separates average density from spatial heterogeneity; PC2 separates high-aggressiveness, high-viability tumors from low-viability, smaller tumors. (B) Hierarchical clustering dendrogram (Euclidean distance, Ward’s linkage) of 18 variables in FAMD space, identifying three clusters: Cluster 1 (raw levitation averages + Ki67 + age), Cluster 2 (heterogeneity metrics), and Cluster 3 (xmc_avg + clinical staging + viability). Horizontal dashed line denotes the cutoff height (1.113) used to define cluster membership. (C) Hierarchical clustering using cosine distance (cutoff height: 0.758), yielding a convergent three-cluster structure that further confirms integration of xmc with staging variables across both distance metrics. Variable color coding as in (A).

Hierarchical clustering using Euclidean distance identified three variable clusters (Figure 3B; Supplementary Table S8): Cluster 1 (n=5) grouped raw levitation averages with Ki67 and age; Cluster 2 (n=3) isolated heterogeneity metrics into an orthogonal spatial variability group; Cluster 3 (n=10) co-clustered xmc_avg with staging variables (T, N, molecular and histological subtype), aggressiveness, and viability. The integration of xmc with clinical staging in Cluster 3 is notable despite only moderate pairwise distances (xmc to N: 0.54; xmc to T: 0.55), suggesting shared latent biological factors beyond simple linear correlations. Complementary clustering using cosine distance yielded a convergent three-cluster structure (Figure 3C; Supplementary Table S9) in which both distance metrics produced mixed levitation-clinical clusters with no purely clinical group emerging, providing convergent evidence that levitation captures information fundamentally intertwined with breast cancer biology.

### Latent Biological Axes Identified by PLS Regression

To identify shared latent factors underlying the FAMD co-clustering, we performed exploratory dual-outcome PLS Regression (PLS-R) with xmc_avg and Heterogeneity_xmc as outcomes (Y-block) and 13 clinical/viability variables as predictors (X-block; n=17 patients; Figure 4A). The two-component model captured moderate variance (R²=0.33) but showed poor cross-validation (Q²=−0.20) and non-significant permutation testing (p=0.63; Supplementary Table S10). These metrics classify the model as exploratory and hypothesis-generating rather than predictive; the biological structure revealed by latent variable loadings is intended to guide validation in larger cohorts.

**Figure 4.**
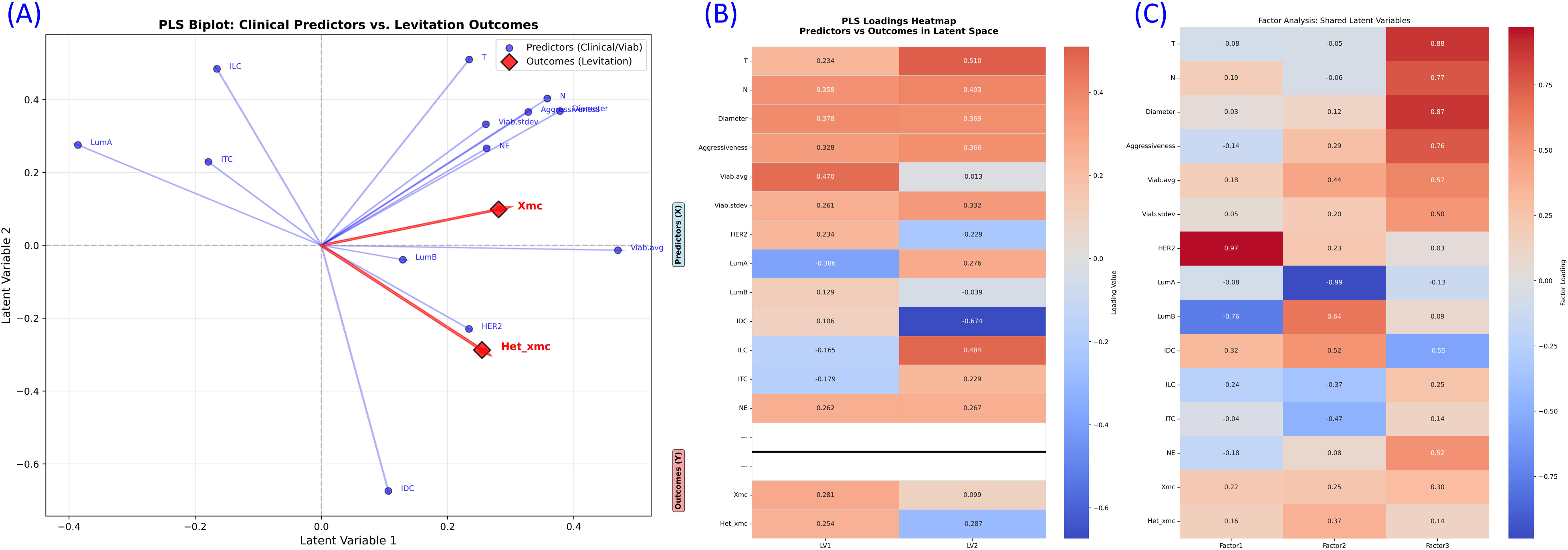
Latent variable structure of levitation–clinical covariance identified by PLS regression and factor analysis. (A) PLS biplot showing the projections of clinical/viability predictors (blue) and levitation outcomes (xmc_avg and Heterogeneity_xmc; red) onto LV1 and LV2. The distance from the origin reflects each variable’s contribution to the model; the angle between vectors reflects their correlation within the latent space. (B) Loading heatmap for the dual-outcome PLS-R model. Rows correspond to X-block predictors (clinical and viability variables, upper block) and Y-block outcomes (xmc_avg and Heterogeneity_xmc, lower block, separated by a horizontal divider); columns correspond to LV1 and LV2. Color intensity reflects loading magnitude; red indicates positive loading, blue indicates negative loading. (C) Factor analysis loading heatmap (varimax rotation, 3 factors) for the combined X- and Y-block variable space. Row order matches panel B; columns correspond to Factor 1, Factor 2, and Factor 3. Both levitation outcomes show distributed loadings across all three factors, indicating that magnetic levitation integrates information from multiple biological dimensions.

Latent Variable 1 (24.6% clinical variance, 14.3% levitation variance) captured a viability-burden axis. Variable Importance in Projection (VIP) analysis identified tissue viability as the dominant driver (VIP=1.60, loading=0.47), followed by nodal status (VIP=1.44, loading=0.36), LumA molecular subtype (loading=−0.39), and tumor diameter (loading=0.38; Figure 4B; Supplementary Figure S7). Both levitation outcomes loaded positively on LV1 (xmc_avg: 0.28; Heterogeneity_xmc: 0.25), indicating that metabolically active, node-positive, non-LumA tumors exhibit both denser average profiles and greater spatial heterogeneity. Notably, nodal status ranked second across all clinical variables despite only moderate univariate correlation with levitation metrics (xmc vs. N: r=0.11, p=0.67).

Latent Variable 2 (13.2% clinical variance, 18.5% levitation variance) defined an architectural axis dominated by histological subtype (IDC loading=−0.67; ILC loading=0.48) and tumor size (T-stage: 0.51). Heterogeneity_xmc loaded negatively on LV2 (−0.29) while xmc_avg loaded near zero (0.10), indicating that architectural features modulate spatial heterogeneity independently of average density: large ILC tumors—characterized by single- file infiltrative growth—showed reduced levitation heterogeneity, while IDC tumors with irregular glandular structures and desmoplastic stroma showed the opposite pattern.

Convergent Factor Analysis on the combined variable space (15 variables, 3 factors with varimax rotation) yielded three interpretable dimensions (Figure 4C): a staging-aggressiveness factor (T: 0.88, N: 0.77, Diameter: 0.87, Aggressiveness Score: 0.76), a molecular subtype factor (LumA: −0.99, LumB: 0.64), and a HER2-specific factor (0.97). Both levitation outcomes showed distributed loadings across all three factors (xmc_avg: 0.22–0.30; Heterogeneity_xmc: 0.14–0.37), confirming that magnetic levitation integrates information from multiple biological dimensions rather than aligning with any single clinical axis

### Patient Stratification in Biophysical Latent Space

Projecting all 17 patients into the two-dimensional latent space (Figure 5A) revealed non-random distribution (spread ratio=0.55; Supplementary Table S11), with 41% of patients in the high-LV1/low-LV2 quadrant characteristic of viable, node-positive, IDC-predominant tumors. Unsupervised K-means clustering identified three natural patient subgroups as optimal (silhouette score=0.51 at k=3; Supplementary Table S12). Cluster 1 (n=9, 53%) comprised exclusively IDC cases with mixed LumB (56%) and HER2-enriched (33%) subtypes (Figure 5B). Cluster 2 (n=5, 29%) was predominantly ILC/LumA and showed the lowest metabolic activity (0.19). Cluster 3 (n=3, 18%) occupied extreme high-LV1/high-LV2 space and represented the most aggressive cases (Figure 5B–5C): 289% larger tumor diameter than Cluster 1 (50.0 vs. 17.2 mm; p<0.001), 88% higher viability (0.56 vs. 0.30; p<0.001), and 242% higher nodal burden (mean N=2.67 vs. 1.11; p<0.001). Aggressiveness Score showed a stepwise gradient across clusters (Cluster 2: 2.8 < Cluster 1: 4.2 < Cluster 3: 6.7; p=0.002). Cluster 2’s low metabolic activity is consistent with the low cell number characteristic of LumA and ILC subtypes (32,33). No significant age differences were observed between clusters (p=0.31), confirming that stratification reflects tumor biology rather than demographic confounding. Clinical variable gradients across latent space confirmed the two-axis model (Supplementary Table S13): viability showed the strongest gradient predominantly along LV1 (R²=0.75, βLV1=2.63), validating LV1 as the metabolic-burden axis. T-stage varied orthogonally along LV2 (R²=0.66, βLV2=0.51). Molecular subtypes showed strong spatial segregation (mean inter-category distance=2.69, 1.7× overall patient spread): LumA tumors clustered in negative-LV1/positive-LV2 space (−2.30, 0.88; n=4), HER2-enriched in positive-LV1/negative-LV2 space (+1.39, −0.73; n=4), and LumB at intermediate positions (n=9). This molecular separation, independent of but complementary to histological separation along LV2, demonstrates that magnetic levitation integrates molecular and architectural tumor characteristics into a unified biophysical signature.

**Figure 5.**
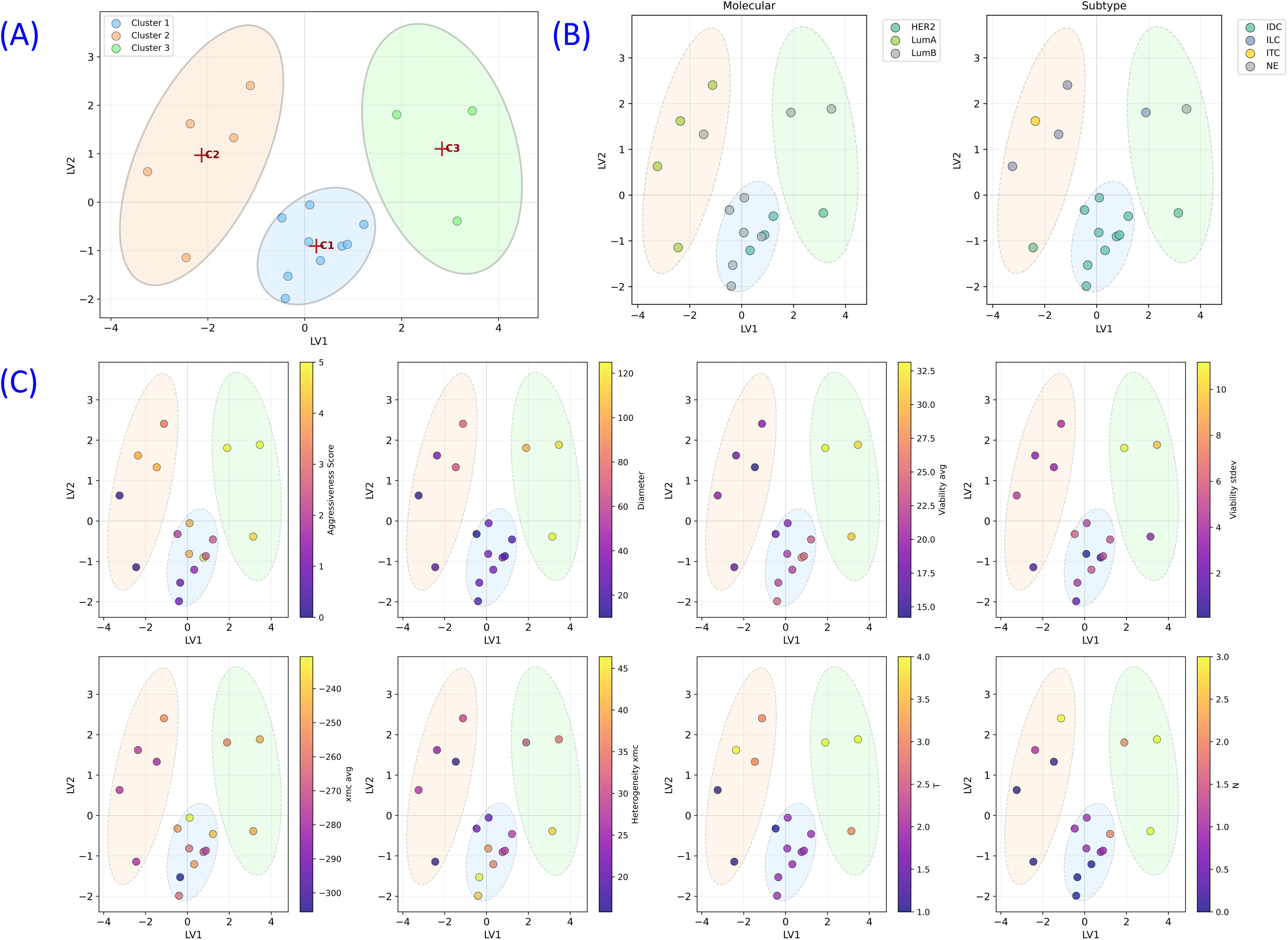
Patient stratification in PLS-derived biophysical latent space. (A) Distribution of all 17 patients in the two-dimensional latent space defined by LV1 (x-axis; viability-burden axis) and LV2 (y-axis; architectural-staging axis), colored by K-means cluster assignment (k = 3, silhouette score = 0.505) with marked centroids. (B) Same latent space axes as (A), patient symbols colored by molecular subtype (left; LumA, LumB, HER2-enriched) and histological subtype (right; IDC, ILC, ITC, NE). (C) Gradient projections of eight continuous variables onto the LV1–LV2 space: Aggressiveness Score, tumor diameter, Viability_avg, Viability_stdev, xmc_avg, Het_xmc, T-stage (T), and nodal status (N). Each panel uses the same LV1/LV2 axes as (A), with patient symbols colored by the respective variable value (low-to-high color gradient).

## DISCUSSION

This study establishes tissue-scale magnetic levitation as a viable biophysical profiling platform for clinical breast cancer specimens, advancing the technology from single-cell applications to intact biopsy analysis. Through evaluation of 203 samples from 17 patients, we address three methodological barriers that have prevented tissue-scale levitation: size-dependent geometric artifacts requiring physics-based and statistical correction, heterogeneity quantification as a predictive signal rather than noise, and multivariate integration to extract clinically relevant information from composite density profiles.

The key departure from prior MagLev paradigms is that spatial heterogeneity—not average density—predicts metabolic viability. In single-cell studies, viability and levitation height are monotonically coupled through membrane integrity (24,25). Intact tissue biopsies, however, integrate cellular, stromal, and architectural components into a composite density profile where this relationship no longer holds. Tissue-scale heterogeneity likely reflects microenvironmental complexity: proliferative zones with high nuclear-to-cytoplasmic ratios (locally denser), quiescent regions with lipid accumulation (less dense), necrotic foci with edema and cell debris (variable density), and collagen-rich desmoplastic stroma (1.3–1.4 g/cm³) intermixed with cellular parenchyma (∼1.05–1.08 g/cm³) (11,34). Metabolically compromised tissues, by contrast, may undergo spatially uniform degradation, producing homogeneous levitation profiles with low heterogeneity despite altered average density. This framework finds a natural analogy similar to that in radiomics, where image texture features quantifying spatial heterogeneity in CT and MRI consistently outperform first-order statistics in predicting tumor biology (35,36). We propose that levitation operates as density-based texture analysis: xmc_avg functions as a first-order feature of bulk tissue composition, while Heterogeneity_xmc serves as a higher-order feature of architectural complexity—a tissue-scale emergent property invisible at single-cell resolution.

The identification of nodal status as the strongest multivariate predictor constitutes our most unexpected finding. Nodal involvement showed only weak univariate associations with levitation metrics but dominated PLS loadings in the context of other clinical variables—a pattern indicating multivariate rather than direct biophysical coupling. We propose three non-mutually exclusive mechanisms. First, clonal selection for metastatic competence may enrich primary tumors for cells with distinct biophysical properties: successful lymphatic dissemination requires altered cytoskeletal organization, membrane cholesterol content, and nuclear architecture (37,38)—all potential determinants of bulk density. Even subclonal enrichment (10–30%) could shift the bulk tissue levitation profile detectably. Second, lymphatic invasion may induce paracrine remodeling of the primary tumor microenvironment through VEGF-C/VEGFR-3 signaling, altering lymphatic vessel density, extracellular matrix composition, and immune infiltration patterns (39,40)—all of which affect tissue density. Third, nodal involvement and density signature may share upstream drivers: highly proliferative tumors with elevated nuclear-to-cytoplasmic ratios and ribosomal content are intrinsically denser and simultaneously more likely to metastasize (41). Our finding that nodal VIP scores exceed those of molecular subtype suggests that biophysical signatures may integrate functional metastatic capacity in ways static genomic classifiers cannot, and warrants prospective evaluation as a biomarker of dissemination potential.

The transition from single-cell to tissue-biopsy scale necessitated the methodological innovations presented here. Our physics-based correction (xmc) addresses the key geometric artifact—above-midpoint tissue extension—by recalculating center-of-gravity for the sub-midpoint tissue fraction alone. This correction distinguished itself uniquely by co-clustering with clinical staging variables in FAMD analysis, a pattern not observed for raw or statistically residualized metrics, suggesting it isolates a biologically informative density component. Compared to label-based microfluidic diagnostics that rely on surface biomarkers (EpCAM, HER2, CEA) and are vulnerable to receptor loss through epithelial-mesenchymal transition (42,43), our label-free approach provides a physiologically relevant density signature independent of molecular tag expression. A notable parallel is the Electrical Pathological Analyzer, an impedance-based sensor validated on 313 samples from 68 patients for rapid intraoperative margin assessment (44); our platform similarly provides rapid stratification in a 30-minute assay from standard core biopsies, with the additional capability of multivariate clinical integration. From a translational standpoint, the platform’s primary advantages are low cost (∼$1 per assay), minimal sample preparation (no fixation, staining, or molecular extraction), and rapid turnaround (∼30 minutes). In low- and middle-income countries, where histopathological grading and immunohistochemistry often represent the only available diagnostic modalities and genomic profiling remains economically prohibitive (45,46), levitation-based profiling could serve as an intermediate-tier stratification tool: high-viability, high-heterogeneity signatures could triage patients for intensified treatment even in the absence of molecular testing. In resource-rich settings, the platform may address specific clinical gaps, including ambiguous intermediate-grade tumors (grade 2; ∼40% of IDCs (47)) where chemotherapy benefit is unclear, serial response monitoring during neoadjuvant therapy, and intraoperative margin assessment as a complement to frozen section analysis (48).

The study has important limitations that define priorities for future validation. The primary limitation is sample size (n=17 patients), yielding a predictor-to-sample ratio below the recommended 3:1 minimum for robust multivariate modeling (49). Consequently, PLS models showed negative cross-validation and non-significant permutation testing, classifying findings as exploratory. However, five independent analytical approaches (Pearson correlations, FAMD with two distance metrics, PLS regression, factor analysis, and latent space clustering), each with different assumptions, yielded consistent biological interpretations (viability-burden and architectural axes, nodal status importance, heterogeneity-viability coupling)—a degree of convergence unlikely to arise from statistical artifact alone. The cohort also lacked TNBC cases, limiting generalizability to a biology known for distinct density-relevant features (high proliferation, basal-like phenotype, metabolic stress) (50). An independent validation cohort with balanced molecular subtype representation is essential for clinical utility claims. Biopsy fragments of 200–600 μm may not fully capture macroscopic tumor heterogeneity, and sampling strategy (peripheral vs. core regions) should be systematically evaluated in future studies. Finally, scale-up to routine clinical practice will require automation of tissue preparation and standardization of levitation protocols across sites (51).

Looking ahead, multiplexed levitation using capillary arrays could enable simultaneous profiling of multiple samples or drug conditions. Integration with microfluidics could support continuous-flow density profiling, and expansion to other solid tumors where tissue biopsy is routine would broaden the platform’s scope. Ultimately, positioning tissue levitation as a biophysical dimension within multimodal machine learning frameworks—alongside radiomics, digital pathology, and spatial transcriptomics—offers the most compelling path toward improved, accessible cancer diagnostics (52). In conclusion, this work demonstrates that macroscopic magnetic levitation is not merely a mechanical measurement but a fundamental biophysical readout of tumor heterogeneity and microenvironmental complexity. By translating density profiling from the single-cell paradigm to tissue-scale architectures, we show that emergent macroscopic characteristics—from lymphatic crosstalk and immune infiltration to metabolic reprogramming and extracellular matrix remodeling—manifest as measurable shifts in bulk tissue levitation profiles. The convergence of analytical evidence presented here provides a compelling biological rationale for prospective clinical validation.

## DECLERATIONS

## Acknowledgments

Authors gratefully acknowledge financial support from Izmir Institute of Technology research grants (2022IYTE-2-0047 and 2022IYTE 2-0030). We gratefully thank H. Cumhur Tekin and Ceyda Oksel-Karakus for helpful and valuable suggestions.

## Author contributions

**MG**: Writing – original draft, Methodology, Investigation, Formal analysis, Conceptualization. **ZEG**: Methodology. **MA**: Investigation, Methodology, Conceptualization. **DP**: Supervision, Methodology, Conceptualization. **BB**: Formal analysis, Investigation. **MG**: Investigation, Methodology, Conceptualization. **OY**: Supervision, Resources, Funding acquisition, Methodology, Conceptualization. **GM**: Writing – original draft, Supervision, Resources, Project administration, Funding acquisition, Methodology, Conceptualization. **EO**: Writing – original draft, Supervision, Resources, Project administration, Funding acquisition, Methodology, Formal analysis, Investigation, Conceptualization.

