## Supplementary Information for "Magnetic Levitation of Intact Tumor Biopsies Reveals Multivariate Biophysical Signatures of Breast Cancer Aggressiveness"

**SUPPLEMENTARY METHODS**

**FAMD and Hierarchical Clustering**

Factor Analysis of Mixed Data (FAMD) was performed using the sklearn library in Python to integrate continuous levitation and viability metrics with categorical clinical variables (n = 17 patients, 18 total variables after removing incomplete cases). FAMD extends principal component analysis to handle mixed data types by applying standard PCA to continuous variables and multiple correspondence analysis (MCA) to categorical variables, then combining both in a unified latent space. Variables with zero loading contributions or missing data were excluded prior to analysis; statistical residual metrics (xm_res, xmin_res) were excluded to avoid multicollinearity with their parent raw metrics.

Variable contributions (loadings) to PC1 and PC2 were extracted to interpret biological dimensions. Hierarchical clustering was performed on the FAMD-transformed variable coordinates using two distance metrics: (**1**) Euclidean distance (magnitude-based) with Ward's linkage, and (**2**) cosine distance (1 − cosine similarity; angle-based, independent of magnitude) with Ward's linkage. Optimal cluster count was determined for each by the elbow method applied to the dendrogram, identifying discontinuities in cluster fusion height (cutoff heights: Euclidean = 1.113; cosine = 0.758). Clustering was implemented using scipy.cluster.hierarchy. Pairwise distances between variables within and across clusters were calculated to assess clustering coherence.

**Partial Least Squares Regression (PLS-R)**

Exploratory dual-outcome PLS-R was conducted using scikit-learn's PLSRegression with two latent variables (LV1, LV2). The X-block consisted of 13 standardized predictors: age, Ki67, tumor diameter, lymph node count, molecular subtype (4 binary dummy variables: LumA, LumB, HER2, TNBC), histological subtype (4 binary dummy variables: IDC, ILC, ITC, NE), T-stage (numeric), N-stage (numeric), Viability_avg, and Viability_stdev. The Y-block contained two correlated levitation outcomes: xmc_avg and Heterogeneity_xmc. All variables were z-scored prior to analysis to ensure equal weighting.

Model performance was assessed via R² (in-sample variance explained in the Y-block), Q² (leave-one-out cross-validation R²), and permutation testing (1,000 permutations; p-value = proportion of permuted models with R² ≥ observed R²). Variable Importance in Projection (VIP) scores were calculated as:

$${VIP}_{j}=\sqrt{p\times\sum_{k} \left[ w_{jk}^{2}\times\frac{{SSY}_{k}}{{SSY}_{total}} \right]}$$

where *p* is the number of predictors, *w*_jk_ is the weight of predictor *j* on component *k*, SS*Y*_k_ is the sum of squares explained by component *k*, and SS*Y*_total_ is the total sum of squares in the Y-block. Predictors with VIP > 1.0 were considered important contributors.

**Convergent Factor Analysis**

Factor analysis was performed on the combined X- and Y-blocks (15 variables total) using principal axis factoring with varimax rotation (n_factors = 3, determined by Kaiser criterion [eigenvalue > 1] and scree plot inspection) via sklearn.decomposition.FactorAnalysis. Loadings ≥ 0.40 were interpreted as meaningful associations.

**Latent Space Projection and Clustering**

Patient LV1 and LV2 scores from PLS-R were used as coordinates in a two-dimensional latent space. Spatial distribution was characterized by: (**1**) centroid (mean LV1, mean LV2 across all patients); (**2**) individual Euclidean distances from the centroid; (**3**) mean distance (spatial spread); and (**4**) spread ratio (mean distance / maximum distance). Unsupervised K-means clustering (sklearn.cluster.KMeans) was applied with k = 2–5 clusters, using Euclidean distance and 100 random initializations per k. Optimal k was selected by maximizing the average silhouette coefficient (sklearn.metrics.silhouette_score). Cluster enrichment for clinical features was assessed by ANOVA (continuous variables) and chi-square tests (categorical variables). Clinical variable gradients across latent space were quantified by fitting bivariate linear models of the form: Clinical_Variable = β₀ + β₁×LV1 + β₂×LV2, reporting spatial R², individual coefficients, and gradient magnitude (√[β₁² + β₂²]). Molecular subtype spatial segregation was assessed by computing pairwise Euclidean distances between subtype centroids and comparing the mean inter-category distance to overall patient spread (separation ratio).

Mahalanobis distances from the group centroid were calculated for each patient to identify potential outliers; no patient exceeded 2σ.

**SUPPLEMENTARY TABLES**

**Supplementary Table S1**. Morphological parameters of levitated biopsy samples (n = 203 samples). Area, maximum cross-sectional area of the segmented tissue outline. Height, maximum vertical span of the tissue within the levitation channel. Width, maximum horizontal span. Circularity, 4π × Area/Perimeter² (1.0 = perfect circle). CV, coefficient of variation.CI_95_L, CI_95_U, 95% confidence interval lower or higher.

| Metric | Mean | Std | CV(%) | CI_95_L | CI_95_U | Median | Q25 | Q75 | Min | Max | Range |
| --- | --- | --- | --- | --- | --- | --- | --- | --- | --- | --- | --- |
| Area(μm²) | 207908 | 58612 | 28 | 199845 | 215971 | 193936 | 161143 | 246947 | 86412 | 380003 | 293591 |
| Perim.(μm) | 3511 | 1061 | 30 | 3365 | 3657 | 3339 | 2741 | 4011 | 1825 | 7581 | 5755 |
| Height(μm) | 499 | 77 | 15 | 489 | 510 | 497 | 446 | 548 | 298 | 752 | 453 |
| Width(μm) | 622 | 132 | 21 | 604 | 640 | 595 | 521 | 715 | 400 | 1017 | 617 |
| Circularity | 0.24 | 0.09 | 38 | 0.23 | 0.25 | 0.23 | 0.17 | 0.30 | 0.07 | 0.49 | 0.42 |

**Supplementary Table S2.** Raw levitation metric distributions (n = 203 samples). All heights are reported relative to the magnetic field midpoint (x = 0 μm); negative values indicate positions below the midpoint. x_m_, center-of-gravity levitation height; x_min_, minimum levitation height (bottom tissue surface); x_max_, maximum levitation height (top tissue surface).

| Metric | Mean(μm) | Std | CV(%) | CI_95_L | CI_95_U | Median | Q25 | Q75 | Min | Max | Range |
| --- | --- | --- | --- | --- | --- | --- | --- | --- | --- | --- | --- |
| xm | -150 | 77 | 51 | -160 | -139 | -135 | -207 | -96 | -327 | 10 | 336 |
| xmin | -399 | 81 | 20 | -410 | -388 | -398 | -466 | -343 | -574 | -227 | 348 |
| xmax | 100 | 91 | 91 | 88 | 113 | 119 | 45 | 162 | -164 | 260 | 424 |

**Supplementary Table S3.** Pearson correlations (r values) between raw levitation metrics and morphological parameters (n = 203 samples). Values without notation are significant at p < 0.05; (ns) = not significant (p ≥ 0.05).

| Metric | Area | Perimeter | Height | Width | Circularity |
| --- | --- | --- | --- | --- | --- |
| xm | 0.243 | 0.150 | 0.149 | 0.241 | -0.101 (ns) |
| xmin | -0.122 (ns) | -0.155 | -0.335 | 0.051 (ns) | 0.077 (ns) |
| xmax | 0.519 | 0.390 | 0.549 | 0.363 | -0.239 |

**Supplementary Table S4.** Univariate correlations between all levitation metrics and mean tissue viability (Viability_avg) at patient level (n = 17). Values are considered statistically significant at p < 0.05; statistical trend at p<0.20; (ns) = not significant (p ≥ 0.20).

| Levitation Metric | Pearson_r | P_value | Difference |
| --- | --- | --- | --- |
| xm_avg | 0.155 | 0.553 | ns |
| xmin_avg | 0.237 | 0.359 | ns |
| xmax_avg | 0.059 | 0.822 | ns |
| xmc_avg | 0.299 | 0.243 | ns |
| xm_res_avg | 0.192 | 0.461 | ns |
| xmin_res_avg | 0.226 | 0.382 | ns |
| Heterogeneity_xm | -0.047 | 0.858 | ns |
| Heterogeneity_xmin | 0.345 | 0.175 | ns |
| Heterogeneity_xmax | -0.263 | 0.307 | ns |
| Heterogeneity_xmc | 0.569 | 0.017 | significant |
| Heterogeneity_xm_res | 0.058 | 0.826 | ns |
| Heterogeneity_xmin_res | 0.297 | 0.247 | ns |

**Supplementary Table S5.** Univariate correlations between levitation heterogeneity metrics and viability dispersion (Viability_stdev) at patient level (n = 17). Values are considered statistically significant at p < 0.05; statistical trend at p<0.20; (ns) = not significant (p ≥ 0.20).

| Levitation Metric | Pearson_r | P_value | Difference |
| --- | --- | --- | --- |
| xm_avg | -0.123 | 0.637 | ns |
| xmin_avg | 0.075 | 0.773 | ns |
| xmax_avg | -0.257 | 0.32 | ns |
| xmc_avg | 0.305 | 0.234 | ns |
| xm_res_avg | 0.031 | 0.905 | ns |
| xmin_res_avg | -0.006 | 0.981 | ns |
| Heterogeneity_xm | 0.41 | 0.102 | trend |
| Heterogeneity_xmin | 0.449 | 0.071 | trend |
| Heterogeneity_xmax | 0.178 | 0.493 | ns |
| Heterogeneity_xmc | 0.008 | 0.976 | ns |
| Heterogeneity_xm_res | 0.466 | 0.059 | trend |
| Heterogeneity_xmin_res | 0.459 | 0.064 | trend |

**Supplementary Table S6.** Multiple linear regression of levitation metrics predicting Viability_avg (AIC-optimized final model; n = 17 patients). Model selection used stepwise backward elimination optimized by AIC, iteratively removing the predictor with the highest p-value until all remaining predictors had p < 0.20.

| **Metric** | | **Value** | | |  |  |  |
| --- | --- | --- | --- | --- | --- | --- | --- |
| Model Type | | Best Model (AIC Optimized) | | |  |  |  |
| Target | | Viability_avg | | |  |  |  |
| R-squared | | 0.911 | | |  |  |  |
| Adjusted R-squared | | 0.764 | | |  |  |  |
| AIC | | 86.55 | | |  |  |  |
| F-statistic P-value | | 0.019 | | |  |  |  |
| N | | 17 | | |  |  |  |
| **Predictor** | **Target_Viability_Metric** | | **Coefficient** | **Std_Error** | | **P_value** | **Significant** |
| xm_avg | Viability_avg | | 5.24E+08 | 1.32E+08 | | 0.007 | Yes |
| xmin_avg | Viability_avg | | 9.92E+08 | 2.5E+08 | | 0.007 | Yes |
| xmax_avg | Viability_avg | | 26462.3 | 7860.991 | | 0.015 | Yes |
| xm_res_avg | Viability_avg | | -5.2E+08 | 1.32E+08 | | 0.007 | Yes |
| xmin_res_avg | Viability_avg | | -9.9E+08 | 2.5E+08 | | 0.007 | Yes |
| Heterogeneity_xm | Viability_avg | | 0.423 | 0.29 | | 0.195 | No |
| Heterogeneity_xmin | Viability_avg | | -1.887 | 0.738 | | 0.043 | Yes |
| Heterogeneity_xmax | Viability_avg | | -0.467 | 0.129 | | 0.011 | Yes |
| Heterogeneity_xmc | Viability_avg | | 0.587 | 0.156 | | 0.009 | Yes |
| Heterogeneity_xmin_res | Viability_avg | | 1.733 | 0.862 | | 0.091 | Trend |

**Supplementary Table S7.** Intraclass correlation coefficients (ICC) of levitation metrics across replicate samples within patients (ICC values reflect the proportion of total metric variance attributable to between-patient differences; the remainder reflects within-patient (intra-tumoral) variability.)

| Levitation Metric | ICC |
| --- | --- |
| xm | 0.178355 |
| xmin | 0.135152 |
| xmax | 0.233892 |
| xmc | 0.151045 |
| xm_res | 0.155737 |
| xmin_res | 0.133465 |

**Supplementary Table S8.** FAMD hierarchical variable clustering — Euclidean distance (magnitude-based). Cutoff height = 1.113 (determined by elbow method on dendrogram fusion heights). **Cluster 1** groups raw levitation averages with proliferative and demographic variables; **Cluster 2** isolates heterogeneity metrics as an orthogonal spatial variability dimension; **Cluster 3** forms a mixed clinical-levitation cluster in which xmc_avg co-groups with staging and aggressiveness variables

| Cluster 1 | Cluster 2 | Cluster 3 |
| --- | --- | --- |
| - xm_avg [Levitation] | - Heterogeneity_xm [Levitation] | - T [Clinical-Cat] |
| - xmin_avg [Levitation] | - Heterogeneity_xmin [Levitation] | - Molecular [Clinical-Cat] |
| - xmax_avg [Levitation] | - Heterogeneity_xmax [Levitation] | - Aggressiveness_Score [Clinical-Cont] |
| - Ki67 [Clinical-Cont] |  | - Diameter [Clinical-Cont] |
| - Age [Clinical-Cont] |  | - Viability_stdev [Viability] |
|  |  | - N [Clinical-Cat] |
|  |  | - Heterogeneity_xmc [Levitation] |
|  |  | - xmc_avg [Levitation] |
|  |  | - Viability_avg [Viability] |
|  |  | - Subtype [Clinical-Cat] |

**Supplementary Table S9.** FAMD hierarchical variable clustering — Cosine distance (angle-based similarity). Cutoff height = 0.758. Cosine distance values range from 0 (identical directional pattern) to 1 (opposing patterns); values ~0.5 indicate unrelated variables.

| Cluster 1 | Cluster 2 | Cluster 3 |
| --- | --- | --- |
| - xm_avg [Levitation] | - Aggressiveness_Score [Clinical-Cont] | - Ki67 [Clinical-Cont] |
| - xmin_avg [Levitation] | - Diameter [Clinical-Cont] | - Heterogeneity_xmax [Levitation] |
| - xmax_avg [Levitation] | - Viability_avg [Viability] | - Heterogeneity_xmin [Levitation] |
| - T [Clinical-Cat] |  | - Heterogeneity_xm [Levitation] |
| - Age [Clinical-Cont] |  | - Molecular [Clinical-Cat] |
|  |  | - xmc_avg [Levitation] |
|  |  | - N [Clinical-Cat] |
|  |  | - Subtype [Clinical-Cat] |

**Supplementary Table S10.** PLS-R model performance summary (dual-outcome model: xmc_avg and Heterogeneity_xmc)

**Overall model performance**

| Metric | Value |
| --- | --- |
| R² (in sample fit) | 0.328 |
| Q² (leave one out CV) | -0.201 |
| Permutation p-value | 0.632 |

**Variance explained by latent variable**

| Block | | | LV1 | LV2 | Total |
| --- | --- | --- | --- | --- | --- |
| X-block (clinical predictors) | | | 24.6% | 13.2% | 37.8% |
| Y-block (levitation outcomes) | | | 14.3% | 18.5% | 32.8% |

**LV1 loadings (viability-burden axis)**

| Latent Variable | Loading | VIP Score |
| --- | --- | --- |
| Viab.avg | 0.470 | 1.603 |
| N | 0.358 | 1.441 |
| Diameter | 0.378 | - |
| LumA | -0.386 | 1.212 |
| xmc_avg (Y) | 0.281 | - |
| Heterogeneity_xmc (Y) | 0.254 | - |

**LV2 loadings (tumor architecture axis)**

| Latent Variable | Loading | VIP Score |
| --- | --- | --- |
| IDC | -0.674 | - |
| T | 0.510 | - |
| ILC | 0.484 | - |
| xmc_avg (Y) | 0.099 | - |
| Heterogeneity_xmc (Y) | -0.287 | - |

**Supplementary Table S11.** Patient latent space distribution summary (n=17). No patient exceeded 2σ Mahalanobis distance from the group centroid, confirming the latent space captures the full phenotypic spectrum without influential outliers driving observed patterns.

| Metric | Value |
| --- | --- |
| Latent Space Dimensions | LV1, LV2 |
| LV1 Range | -3.243 to 3.457 |
| LV2 Range | -1.986 to 2.406 |
| Overall Centroid | LV1= 0.000, LV2= 0.000 |
| Mean Distance from Centroid | 1.959 ± 1.070 |
| Spread Ratio (mean/max distance) | 0.546 |

| Quadrant | Interpretation | Patient Count | Percentage |
| --- | --- | --- | --- |
| Q1: High LV1, High LV2 | Aggressive, high nodal burden | 2 | 11.8 % |
| Q2: Low LV1, High LV2 | Low viability, ILC/LumA | 4 | 23.5 % |
| Q3: Low LV1, Low LV2 | Mixed | 4 | 23.5 % |
| Q4: High LV1, Low LV2 | Viable, node-positive, IDC-predominant | 7 | 41.2 % |

**Supplementary Table S12.** K-means patient cluster analysis in latent space. Optimal number of clusters were 3 according to the Silhouette Scores. Cluster 2's low viability is consistent with the characteristically low proliferative index of LumA and ILC subtypes.

| k | Silhouette Score |
| --- | --- |
| 2 | 0.443 |
| 3 | 0.505 (optimal) |
| 4 | 0.498 |

| Cluster Characteristics | Cluster 1 | Cluster 2 | Cluster 3 |
| --- | --- | --- | --- |
| Patients (%) | 9 (52.9%) | 5 (29.4%) | 3 (17.6%) |
| Centroid (LV1) | 0.239 | -2.129 | 2.833 |
| Centroid (LV2) | -0.905 | 0.969 | 1.101 |
| Intra Cluster Spread | 0.606 | 1.093 | 1.058 |
| Dominant histological subtype | IDC (100%) | ILC 60%, ITC 20% | IDC 67%, NE 33% |
| Dominant molecular subtype | LumB 56%, HER2 33% | LumA 80% | Mixed |
| Mean tumor diameter (mm) | 17.2 | 28.6 | 50.0 |
| Mean Viability_avg | 0.30 | 0.19 | 0.56 |
| Mean N-stage | 1.11 | 0.40 | 2.67 |
| Mean Aggressiveness Score | 4.2 | 2.8 | 6.7 |
| xmc_avg (μm) | −260 | −275 | −252 |
| Heterogeneity_xmc | 32.6 | 22.1 | 35.3 |

**Supplementary Table S13.** Clinical variable gradients across PLS latent space. For each continuous variable, bivariate linear regression was performed: Clinical_Variable = β₀ + β₁×LV1 + β₂×LV2. For categorical variables, mean LV1/LV2 coordinates per category are reported with inter-category distance. Viability aligns predominantly with LV1 (β₁ >> β₂), confirming LV1 as the metabolic-burden axis. T-stage aligns predominantly with LV2 (β₂ >> β₁), confirming LV2 as the architectural-staging axis. Heterogeneity_xmc shows opposing LV1/LV2 coefficients (+/−), indicating bidirectional modulation: increasing with viability/nodal burden along LV1 but decreasing with tumor size/ILC architecture along LV2.

***Continuous variables***

| Variable | LV1 coefficient | LV2 coefficient | Spatial R² | Gradient magnitude |
| --- | --- | --- | --- | --- |
| Viability_avg | +2.629 | −0.074 | 0.752 | 2.630 |
| N_numeric | +0.398 | +0.448 | 0.732 | 0.599 |
| Diameter (mm) | +13.56 | +13.22 | 0.734 | 18.93 |
| T_numeric | +0.233 | +0.508 | 0.660 | 0.559 |
| Aggressiveness_Score | +0.566 | +0.633 | 0.610 | 0.849 |
| xmc_avg (μm) | +5.191 | +1.826 | 0.286 | 5.503 |
| Heterogeneity_xmc | +2.308 | −2.604 | 0.370 | 3.480 |

***Categorical variables — molecular subtype centroids***

| Subtype | n | LV1 centroid | LV2 centroid | Intra-category spread |
| --- | --- | --- | --- | --- |
| LumA | 4 | −2.295 | +0.878 | 1.204 |
| LumB | 9 | +0.402 | −0.066 | 1.453 |
| HER2-enriched | 4 | +1.391 | −0.730 | 0.804 |

Mean inter-category distance: 2.69 (separation ratio: 1.685 × overall patient spread). LumA–HER2 inter-centroid distance: 3.76.

***Categorical variables — histological subtype centroids***

| Subtype | n | LV1 centroid | LV2 centroid |
| --- | --- | --- | --- |
| IDC | 11 | +0.259 | −0.880 |
| ILC | 4 | −0.984 | +1.544 |
| ITC | 1 | −2.364 | +1.620 |
| NE | 1 | +3.457 | +1.884 |

Mean inter-category distance: 3.706 (separation ratio: 2.321 × overall patient spread).

**SUPPLEMENTARY FIGURES**

**Supplementary Figure S1.** Depiction of levitation metrics from optical images. Levitation height metrics xm (center-of-gravity), xmin (bottom surface), and xmax (top surface) are defined relative to the magnetic field midpoint (x = 0 μm)

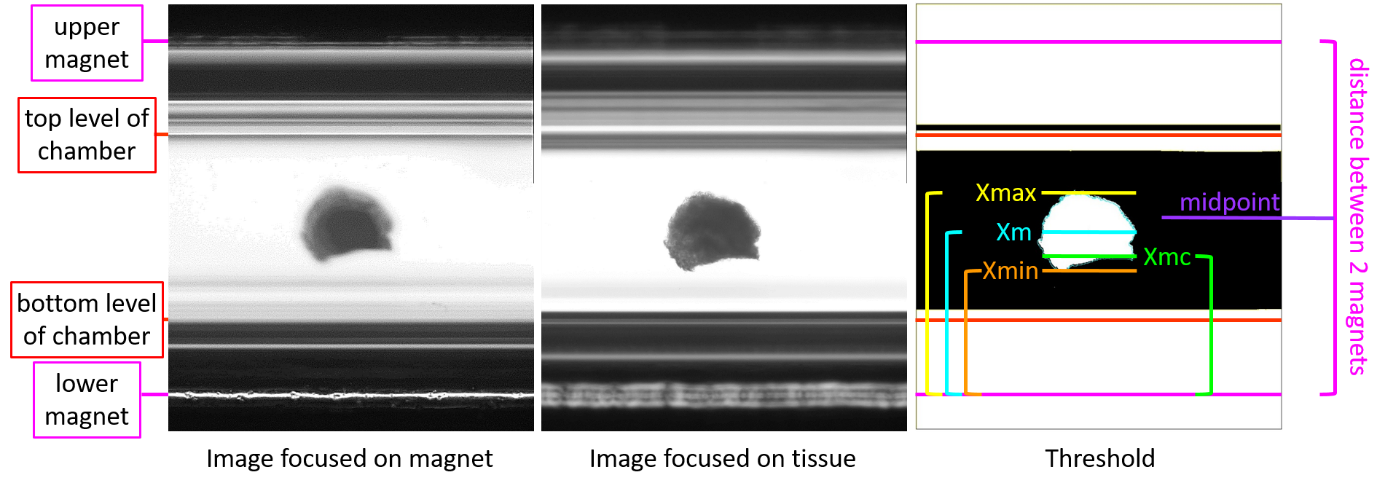

**Supplementary Figure S2.** Size-dependency analysis and normalization comparison. Scatter plots of raw levitation metrics (xm, xmin) versus tissue Area (μm²), with Pearson r and p-values annotated.

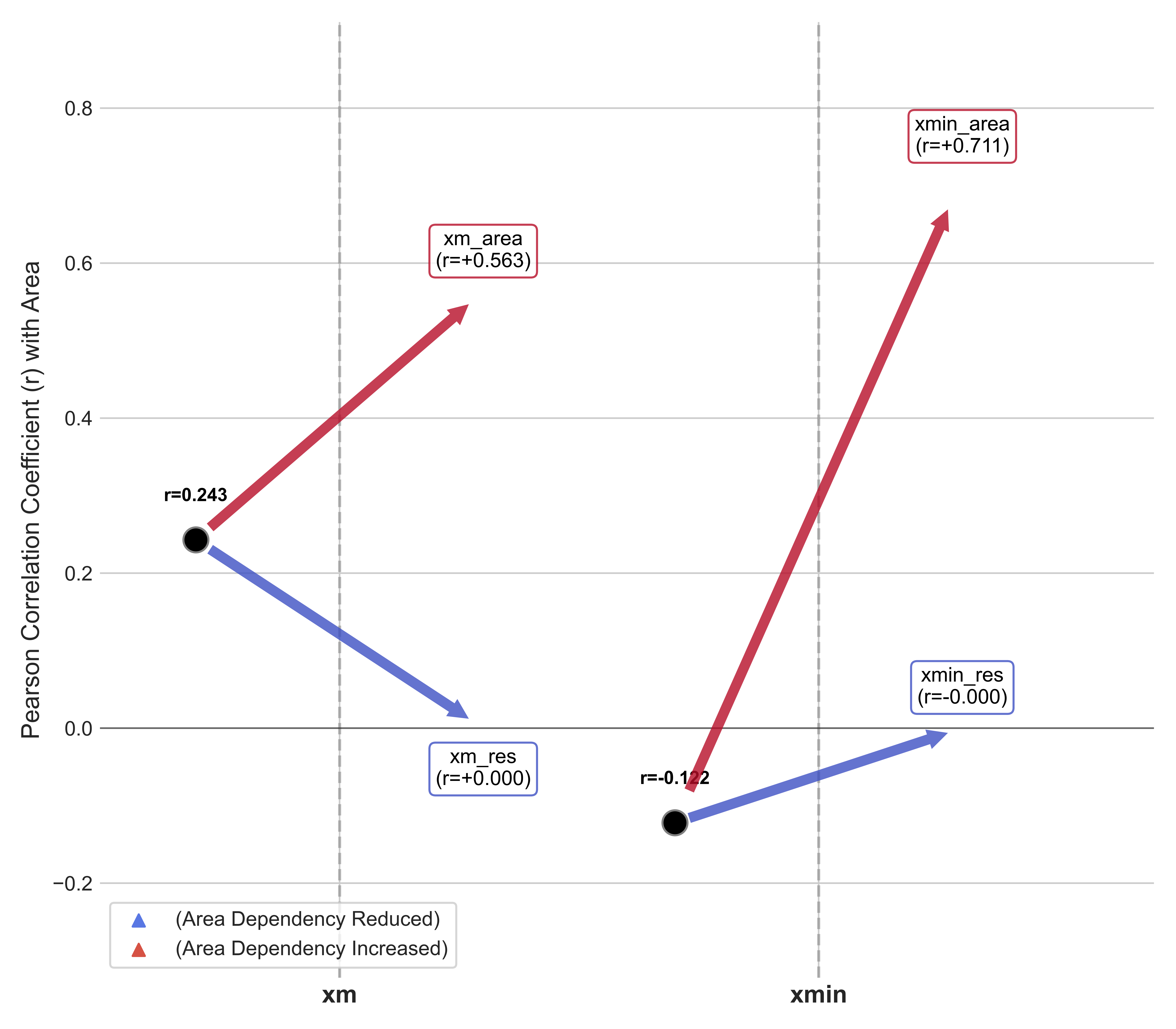

**Supplementary Figure S3.** Heatmap of levitation metric–viability correlations (patient level, n = 17). Pearson r values for all pairwise combinations of 12 levitation metrics (6 averages, 6 heterogeneity measures) against 2 viability metrics (Viability_avg, Viability_stdev)

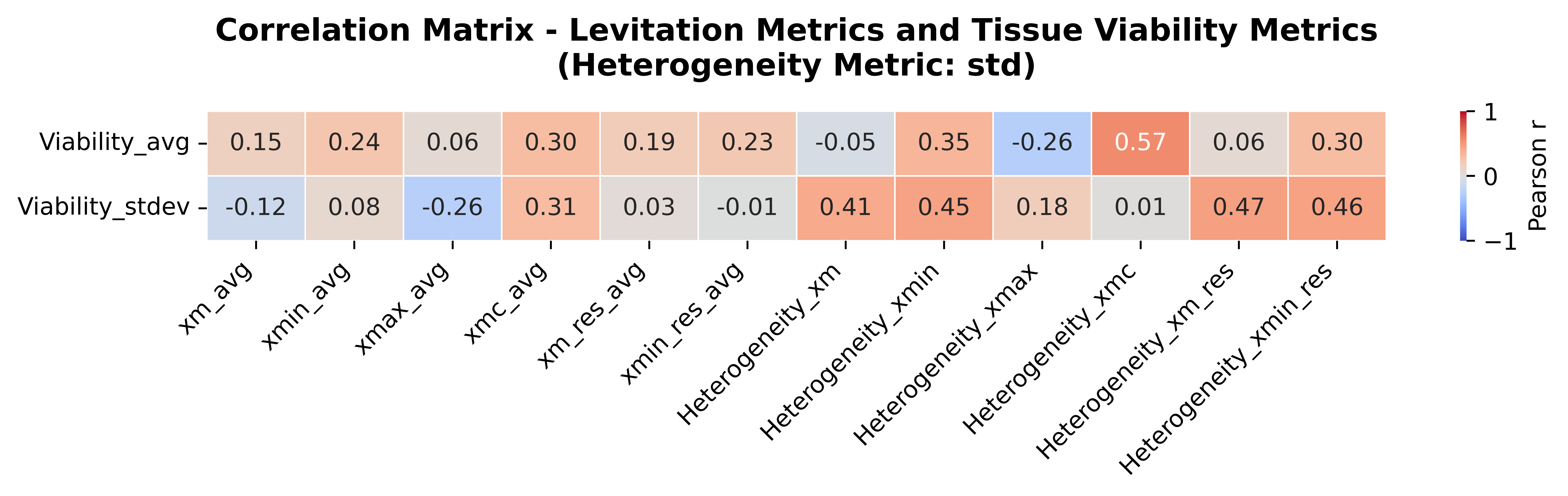

**Supplementary Figure S4.** Patient-level clinical associations (n = 17 patients). Plots depicting associations between patient-averaged viability / levitation metrics and clinical variables. Differences are considered statistically significant at p < 0.05 and showing statistical trend at p<0.20. Horizontal red lines represent mean values, with whiskers corresponding standard deviations.

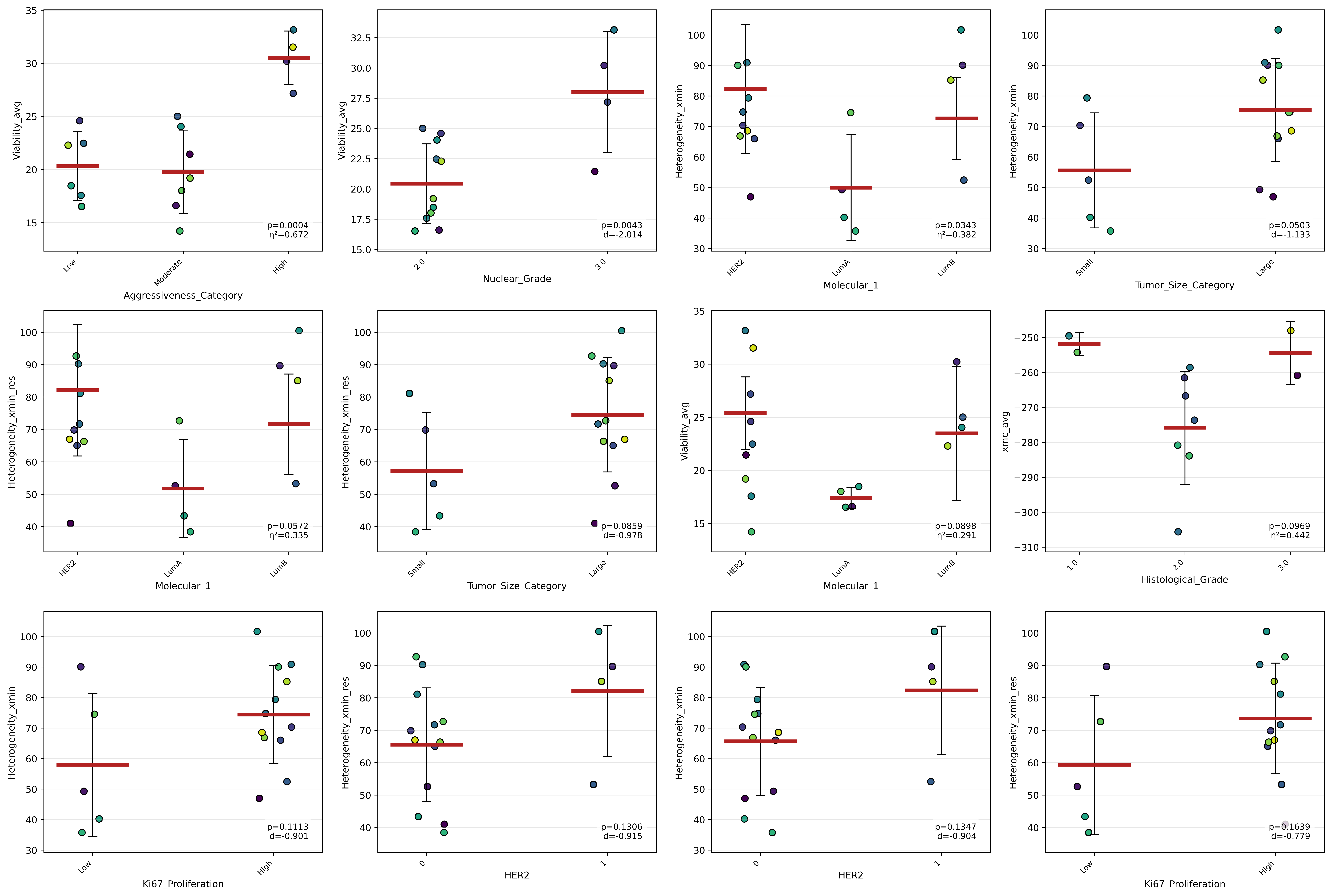

**Supplementary Figure S5.** FAMD scree plot. Variance explained (%) per principal component (PC1–PC10) from the FAMD analysis (n = 17 patients, 18 variables). PC1 and PC2 together explain 40.8% of total variance (PC1: 23.9%; PC2: 16.9%). The dashed vertical line marks the elbow point used to guide interpretation focus on the first two components. The cumulative variance curve (right y-axis) reaches 94.9% by PC10.

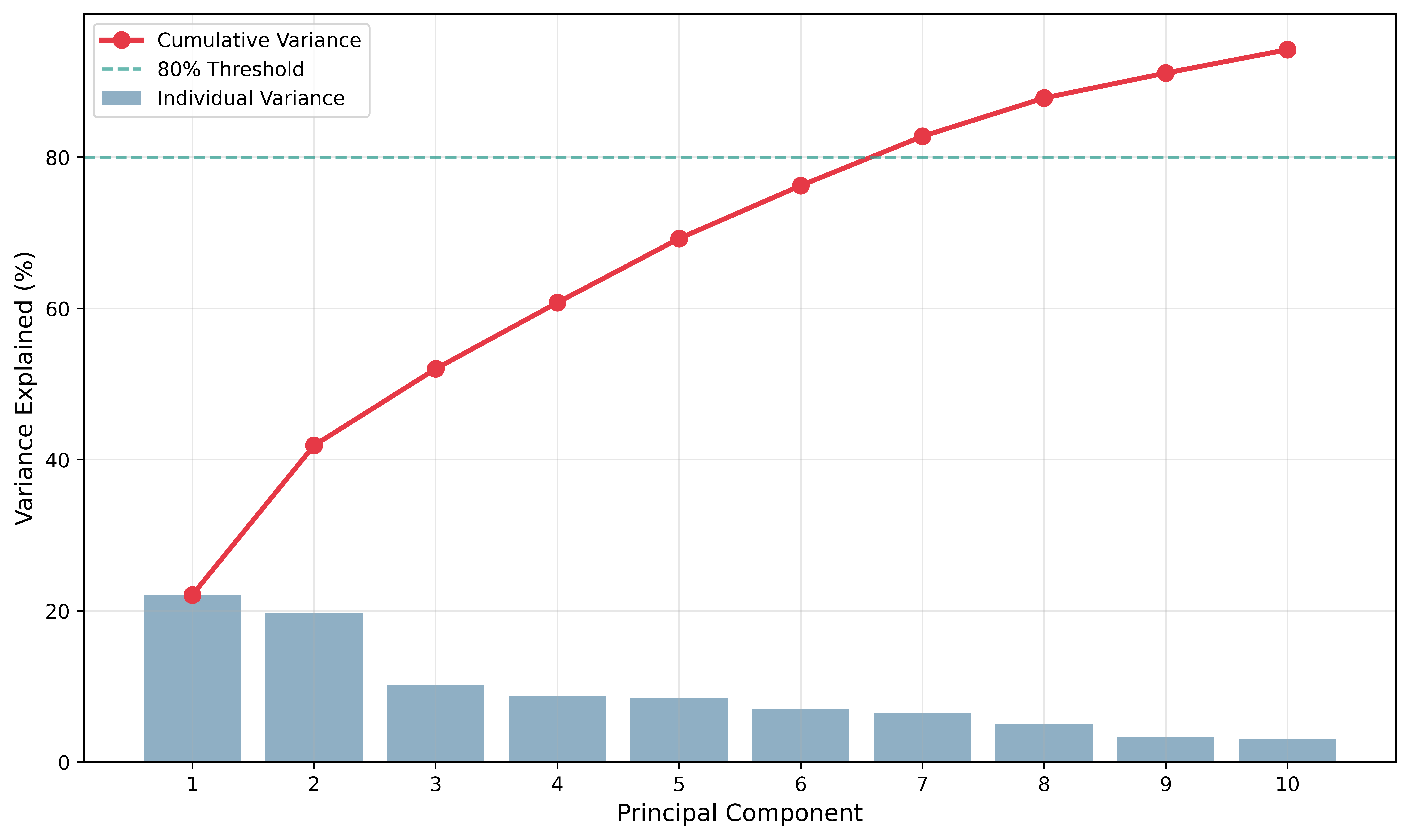

**Supplementary Figure S6.** FAMD variable loadings of all 18 variables on PC1 – PC10.

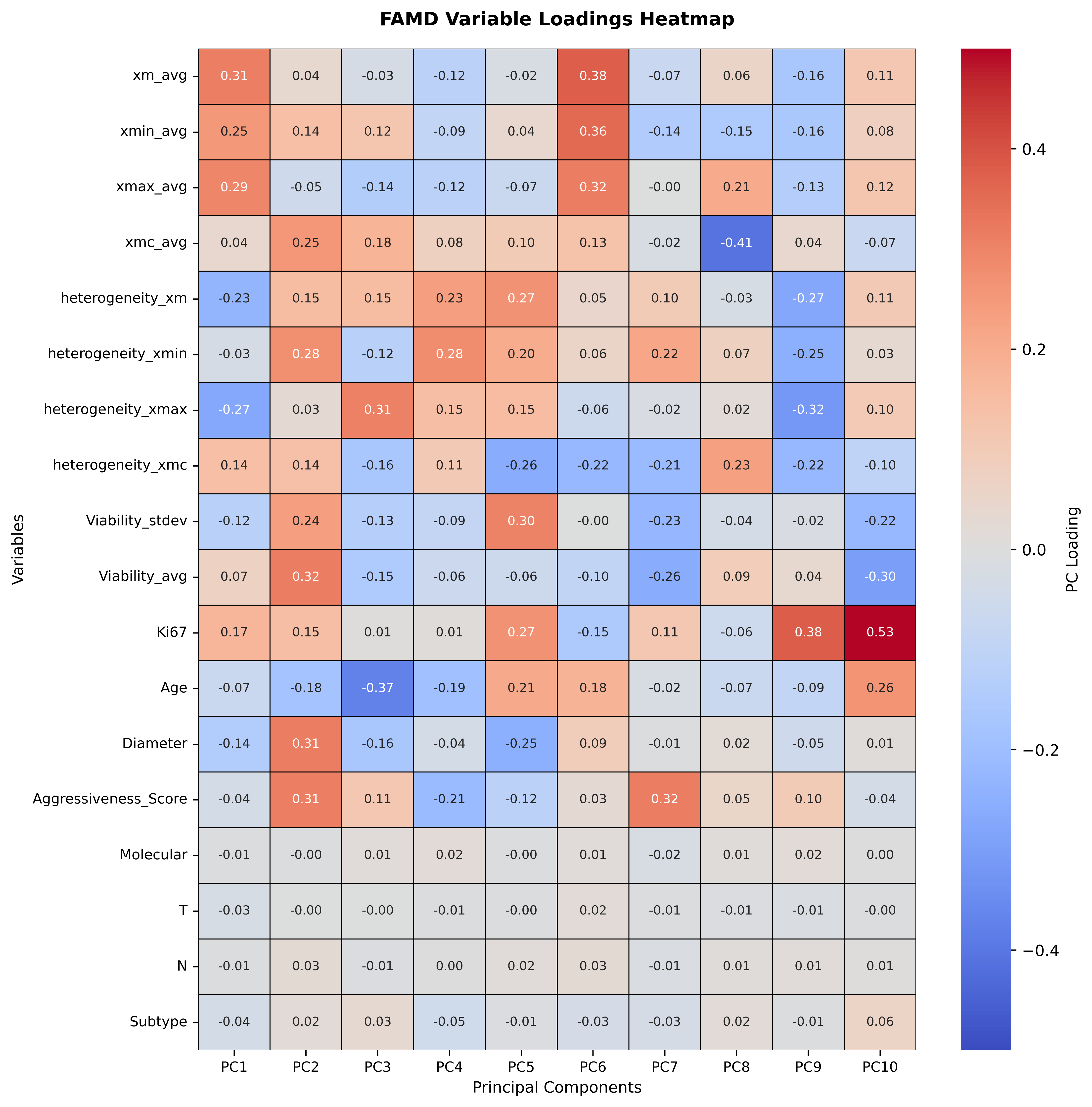

**Supplementary Figure S7.** Variable Importance in Projection (VIP) scores for the dual-outcome PLS-R model. VIP scores for all 13 X-block predictors, ordered by magnitude contributing to higher xmc_avg and greater Heterogeneity_xmc. The dashed horizontal line marks VIP = 1.2 and 1.0.

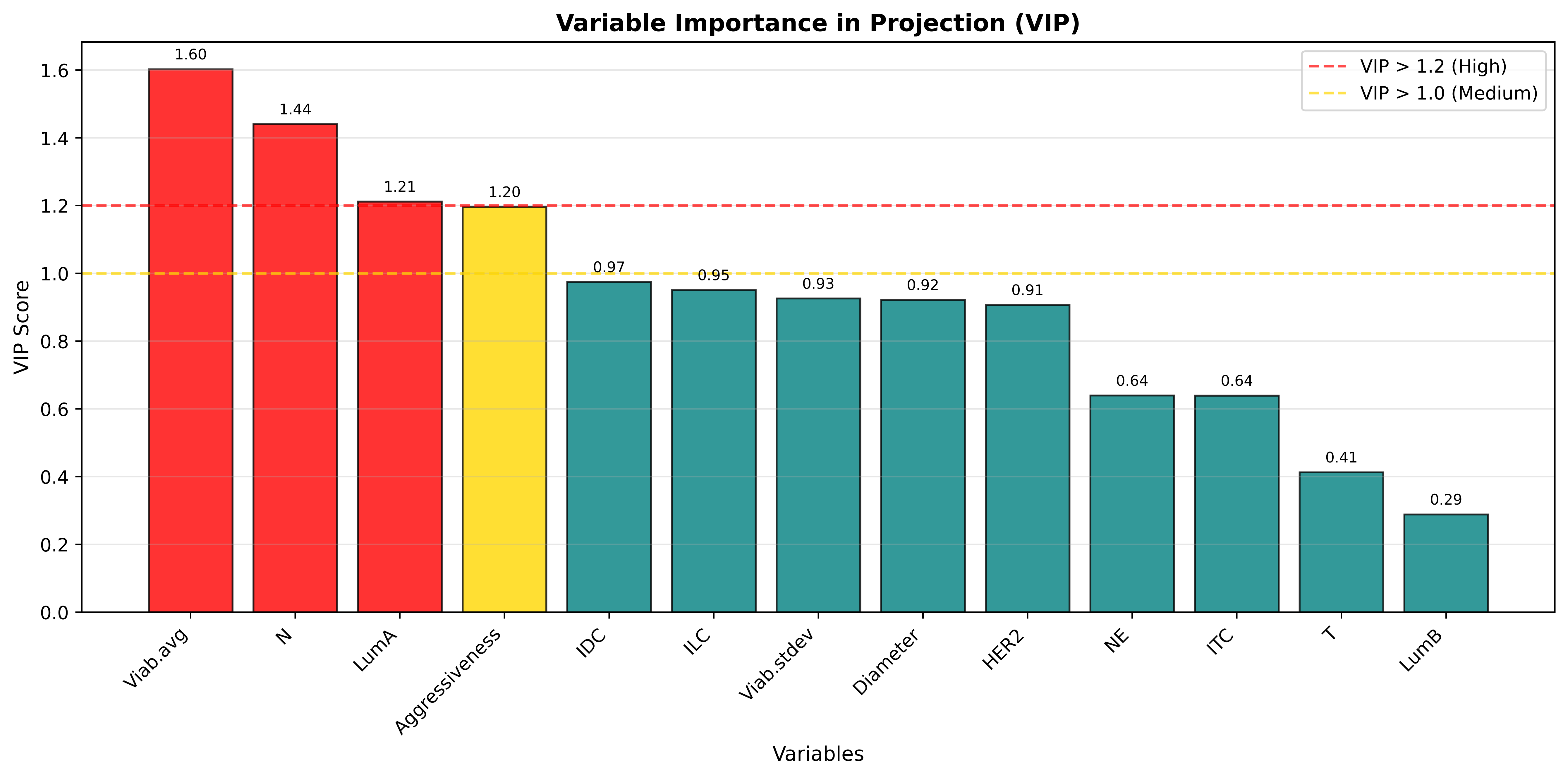
